# PtRNAdb: A web resource of Plant tRNA genes from a wide range of plant species

**DOI:** 10.1101/2022.03.03.482782

**Authors:** Durdam Das, Shafaque Zahra, Ajeet Singh, Shailesh Kumar

## Abstract

tRNA, as well as their derived products such as short interspersed nuclear elements (SINEs), pseudogenes and transfer-RNA, derived fragments (tRFs) has now been shown to be vital for cellular life, functioning and adaptation during different stress conditions in all diverse life forms. In this study, we have developed PtRNAdb (*www.nipgr.ac.in/PtRNAdb*), a plant exclusive tRNA database containing 113849 tRNA gene sequences from phylogenetically diverse plant species. We have analysed a total of 106 nuclear, 89 plastidial and 38 mitochondrial genomes of plants by tRNAscan-SE software package, and after careful curation of the output data, we developed this database and integrated the data. The information about the tRNA gene sequences obtained, were further enriched with consensus sequence based study of tRNA genes based on their isoacceptors and isodecoders. We have also built covariance models based on the isoacceptors and isodecoders of all the tRNA sequences using infernal tool. The user can also perform BLAST not only against PtRNAdb entries but also against all the tRNA sequences stored in PlantRNA databases; and annotated tRNA genes across the plant kingdom available at NCBI. For the users’ ease, we have also incorporated the tRNAscan-SE tool for tRNA gene prediction, and ViennaRNA package for structural analysis on the home page of PtRNAdb. This resource is believed to be of high utility for plant researchers as well as molecular biologists to carry out further exploration of plant tRNAome on a wider spectrum, as well as for performing comparative and evolutionary studies related to tRNAs and their derivatives across all domains of life.

**Database URL:** http://www.nipgr.ac.in/PtRNAdb/

## Introduction

The classical transfer RNA molecules or tRNAs are the most abundant and highly conserved class of non-coding RNAs existing in all life forms. tRNAs with their length ranging from approximately 70-100 nucleotides (nt) can be represented as a cloverleaf with four stems, and the three-dimensional structure as an “L” shape. These distinctly structured molecules act as a connecting bridge between the genetic code and the corresponding translation product (protein) and can be said to be genetic decoders of the code1. Apart from their conventional role as adapters in protein translation, tRNAs play a diverse role in cellular functioning2. In recent years, burgeons of studies related to extra-translational roles of tRNAs have been performed, revealing new dimensions and perspective to the tRNA biology. tRNA can now be regarded as a multi-functional molecular entity governing different aspects of cellular physiology and metabolism in both prokaryotes and eukaryotes alike3. Few identified extra translational functions of both charged as well as uncharged tRNAs in association with cell signalling, stress regulation, and ribosomal stability have been established4,5. These are also involved in the apoptotic regulation process in the mammalian system6. tRNAs upon endonucleolytic cleavage give rise to tRNA halves as well as transfer RNA derived fragments (e.g. tRFs) which are associated with pathogenesis and stress signalling along with other functions in a wide range of organism including plants7–13. tRNA related short interspersed elements (SINES) with unknown functions are also detected in various organisms14–16. The existence of isoacceptors and isodecoders add another layer of intricacy in the tRNA world17,18. Although their most studied roles revolve around protein translation, innumerable complexities pertaining to tRNA biogenesis, processing, intron sequences, pseudogenes, suppressor tRNAs, modifications, interactions with different proteins, diversity and the differential expression of different isoacceptors and isodecoders highlight another level of complexity in the tRNA world19–24. In plants, translation occurs in the cytosol but also in mitochondria and chloroplast25. Thus, there is a higher level of complexity associated with tRNA synthesis, enzymatic machinery involved, localization, and their intricate interactions and functioning within the different compartments of the plant cell26. The gain and loss of introns amongst diverse plant species reveal their significance in the during various stages of evolution 21,27 and aid in studying the comparative genomics across the plant kingdom. The current era with robust Next Generation Sequencing (NGS) technology provides massive opportunity to invade and explore the genomics and extensive transcriptomic repertoire. The sphere of tRNA biology with respect to their structural and functional complications of tRNAs has captivated the attention of scientists from a long time and is continuously flourishing. In order to explore the tRNAomic world in a systematic and specific way from genetics and functional genomics perspective, a number of web repositories have been developed from time to time. Some of the famous and currently available web portals for tRNA related information includes Rfam28, tRNAdb29, GtRNAdb30, tRNADB-CE29 and plantRNA31. The first plant-exclusive web repository, plantRNA was developed in 2013. This knowledgebase harbours annotated tRNA gene sequences and their functions from complete nuclear and organellar genomes from only eleven evolutionarily diverse plants. Due to the availability of a large number of sequenced genomes, there is still a lacuna which requires to be filled for the exploration of vast plant tRNAome in a wider spectrum. Here, we have introduced PtRNAdb (www.nipgr.ac.in/PtRNAdb), the plant exclusive web repository that stores tRNA gene sequences and other related information from 3577 different nuclear as well as mitochondrial, chloroplastic, and plastidial genomes currently available at National Center for Biotechnology Information (NCBI). In addition to the information generated by tRNAscan-SE software package, the intron sequences with their position, length, and GC% have also been incorporated within this database. Additionally, secondary structures of both precursor and mature tRNAs can be also visualized at the search result page. As the consensus transcriptional internal control regions termed as A and B box sequence elements are highly conserved among plants 32,33, are AT-rich34 and, their position in tRNA, as well as CAA motifs, have also been highlighted. CAA motifs are commonly present in the upstream of tRNA genes35 and can act as transcription initiation site. To the best of our knowledge, PtRNAdb is the best platform for plant tRNA genes. The overall representation and major sections of PtRNAdb are shown in Figure 1.

**Figure 1:**
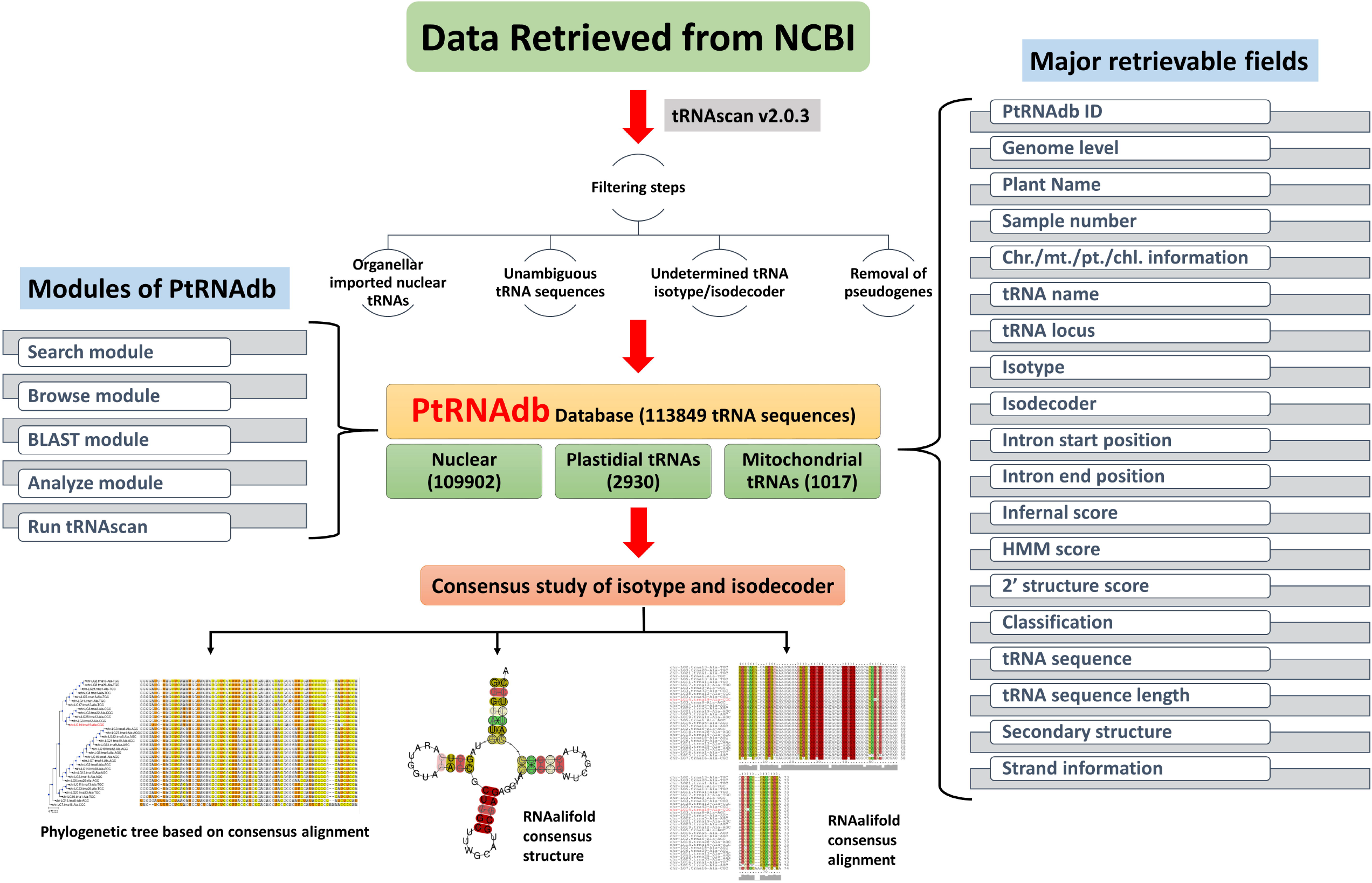
Overview of the web interface, basic functions and features of PtRNAdb.

## Methodology

### Data retrieval

The data was retrieved from National Center for Biotechnology Information (NCBI) database genome browser. The genomes having chromosomal level information were only downloaded and the plant genomes which had scaffold or contigs level of information were excluded to avoid unambiguity in the study. The organellar genomes of only those plants were retrieved for which nuclear data at chromosomal level was also available. The nuclear genome sequence files were downloaded by using File Transfer Protocol (FTP). Further, any extra sequences present in nuclear genomes were manually removed. To download the organellar genomes i.e. mitochondrial and plastidial, the ‘esearch’ and ‘efetch’ utility of NCBI was employed. The ‘E-utilities’ use a fixed URL syntax that translates a standard set of input parameters into the values necessary for various NCBI software components to search and retrieve the requested data.

### tRNA gene prediction

tRNAscan-SE tool developed by Low and Eddy, 1997 (1) was used to predict the tRNA sequences from the genome files. It offers high accuracy and reliable predictions and has been utilized for tRNA gene prediction in tRNA research (2). In this study, tRNAscan-SE (Version 2.0.3) was used with eukaryotic (-E) and organellar (-O) modes for nuclear and organellar genomes respectively. The complete command line options that were used are discussed in the method section of PtRNAdb database (http://14.139.61.8/PtRNAdb/pages/method.php). The output of tRNAscan-SE was saved in six different forms of output files namely (bed, fasta, iso, out, report, ss, stats). Further intensive filtering and analysis was performed on the output data so that only reliable tRNA genes can be presented in the database.

### tRNA sequence filtration

With the aim the to report only reliable and accurate tRNA sequence predictions, various filters were applied on the tRNAscan-SE output. The filtration of tRNAs from the nuclear genomes was done by filtering out the pseudogenes, undetermined tRNAs (i.e. having unambiguous isoacceptors and isodecoders), removing tRNA sequences with more than 2 % of unambiguous nucleotide composition (i.e. N/n), removing large intronic sequences from tRNAs and removing organellar (i.e. mitochondrial and plastidial) imported nuclear tRNAs. In the organellar tRNAs, the undetermined tRNA sequences (i.e. having unambiguous isoacceptors and isodecoders) were filtered out and the tRNA sequences with more than 2 % of unambiguous nucleotide composition (i.e. N/n) were also discarded. After these filtering steps, the quantity of tRNA sequences were reduced but the quality of the overall dataset improved significantly which is proved by further analysis.

### tRNA consensus sequence based study

Followed by the filtering steps, consensus sequence based study was performed on the tRNA sequences of each plant. The consensus study was performed on the isoaccpetors and isodecoders of each plant and the output was a consensus alignment and consensus structure using RNAalifold (3). For this consensus sequence based study, the tRNA sequences from each amino acid was grouped into a single fasta file and similarly, tRNA sequences from each anticodon/isodecoder was collected into a single fasta file. Therefore, as a result of this grouping, now for each plant species there are 20 fasta files according to the isoacceptors and 64 isodecoder wise fasta files. After this, all the fasta files were provided as an input to the LocARNA (4) software for consensus sequence study via an in-house shell script. LocARNA uses Clustal W (5) to align the tRNA sequences. The consensus alignment and the consensus based tRNA structure is available for all the plants present in PtRNAdb and can be accessed either isoacceptors or isodecoder wise from the detailed page of the database. Moreover, a consensus based phylogenetic tree was constructed for each isoacceptor and isodecoder of all plants using Environment for Tree Exploration (ETE) (6). This consensus based phylogenetic tree is also available for all the plants along with the consensus alignment and structure. However, the isoacceptor and isodecoder groups which contain only one tRNA sequence, do not have any consensus based alignment or any consensus based structure since it is only a single sequence. For those single tRNA sequence entries, RNAfold was used which is a utility of ViennaRNA package 2.0 (7) to model the tRNA structure.

### Building tRNA infernal covariance models

Apart from this, probabilistic models were built for each amino acid and anticodon group using various infernal (8) utilities. Probabilistic profiles of the sequence and secondary structure of the isoacceptors and isodecoders was built known as covariance models (CMs). Infernal (“INFERence of RNA ALignment”) (8) is for searching DNA sequence databases for RNA structure and sequence similarities. It is an implementation of a special case of profile stochastic context-free grammars called *covariance models* (CMs). A CM is like a sequence profile, but it scores a combination of sequence consensus and RNA secondary structure consensus, so in many cases, it is more capable of identifying RNA homologs that conserve their secondary structure more than their primary sequence. In order to build these infernal models, firstly, the Stockholm files of each isoacceptor and isodecoder group was provided to “cmbuild” utility of infernal. The Stockholm files were obtained from the previous step in which the consensus study was performed using “mlocarna”. The “cmbuild” utility builds the initial covariance model from an input multiple alignment or Stockholm file. After this, “cmcalibrate” utility is used which calibrates the E-value parameters for the covariance model. In this last step, we need to integrate our models into the database, therefore, “cmpress” utility was used to format a CM database into a binary format for “cmscan” that is integrated in the PtRNAdb. Users can provide their query sequence in the “ANALYZE” module of PtRNAdb database and select the respective plant and isodecoder or isoacceptors model to run cmscan on the query sequence. The output can be downloaded in the result page of Infernal module.

### PtRNAdb web interface development

We have developed PtRNAdb using Apache HTTP server (version 2.4.6) integrated with PHP (version 7.3.3) and MySQL (version 8.0.15) on server machine with Centos 7 Linux as operating system. PHP and JavaScript (version 1.8.0) were used to develop the back-end of the database while MySQL (version 8.0.15) was used to process the data at the back-end. The graphical representation of an overall architecture of PtRNAdb along with the features are displayed in Figure 1. CSS and HTML was used to make the template responsive. Perl and shell scripts were also integrated at the back-end of the database for multiple file handling and data manipulation.

### Features incorporated in PtRNAdb

This database is developed with the aim to provide a user-friendly and simple interface. In order to accomplish this aim, “Search” and “Browse” options are provided on the database. On the search page of PtRNAdb, user can make complex search queries by selecting multiple options and refining the number of output to very specific results or else select less options for broad results. The browse module provides two options to browse through all the entries present in the database: “Browse by tRNA sequence length range” and “Browse by plant family”. This module is very useful in case user do not have any pre-handed information about tRNAs or the database and just wants to understand the features of the database. Apart from the “search” and “browse” modules, “BLAST” module was also incorporated. In this module, we have integrated blastn (9) option from the BLAST software package. This module is very useful to find regions of similarity between the user input FASTA sequences and PtRNAdb database sequences or PlantRNA database sequences or NCBI reported tRNA sequences. User can also manipulate the E-value by using the drop down option on the “BLAST” module which is set at 10 by default.

Apart from the modules, the output tables of the search and browse modules are highly dynamic. In the output search table, the users can further refine the output according to their requirements by using the search box and also shuffle through large number of output entries easily by using the pagination links. It expected that these additional features will make the experience of the user more pleasant and easy to understand the database.

## Results and Discussion

In total, 109902, tRNA genes were registered from the nuclear genome of 106 photosynthetic organisms. We also analysed 127 organellar genomes out of which, 89 samples were plastid genomes and 38 mitochondrial genomes. In the 89 plastid genome samples, we identified 2930 tRNA genes and in 38 mitochondrial genome samples we identified 1017 tRNA genes. Majority of the nuclear as well as organellar genome samples are from dicots i.e. 74 and 91 dicots of nuclear and organellar genomes followed by 29 monocots from nuclear genome and 30 monocots from organellar. Both algae and bryophyte samples were present in low numbers but, no pteridophytes was not present in this study.

The total number of tRNA entries available in the current version of PtRNAdb is 113849. In order to show the relation between the genome size of the plants and the number of tRNAs, pseudogenes and introns, all these data was shown in figure 2 for all the three levels of genomes (i.e. nuclear, plastid and mitochondrial). In part A of figure 2, the plants with nuclear genome is shown and it is quite clear that, with the increase in the genome size, the number of tRNAs, number of pseudogenes and number of introns are increasing. Similarly, in part B and part C of figure 2, a similar graph is visible except there is not pseudogenes. This is because in the tRNAscan-SE algorithm, there is no pseudogene prediction for organellar genome data. Therefore, only genome size (in Kb), number of tRNAs and number of pseudogenes is shown for plastid and mitochondrial genomes. The initial number of tRNAs and the number of true tRNAs obtained after filtration steps is available supplementary table 1. As a result of the filtration steps performed, the quality of the tRNA sequences improved which was found in the consensus sequence study using locARNA. All the results and data available on the database can be retrieved very easily and for understanding each module of the database in a comprehensive manner, user should refer to the help page of PtRNAdb where the usability of all the modules are explained in detail.

**Figure 2:**
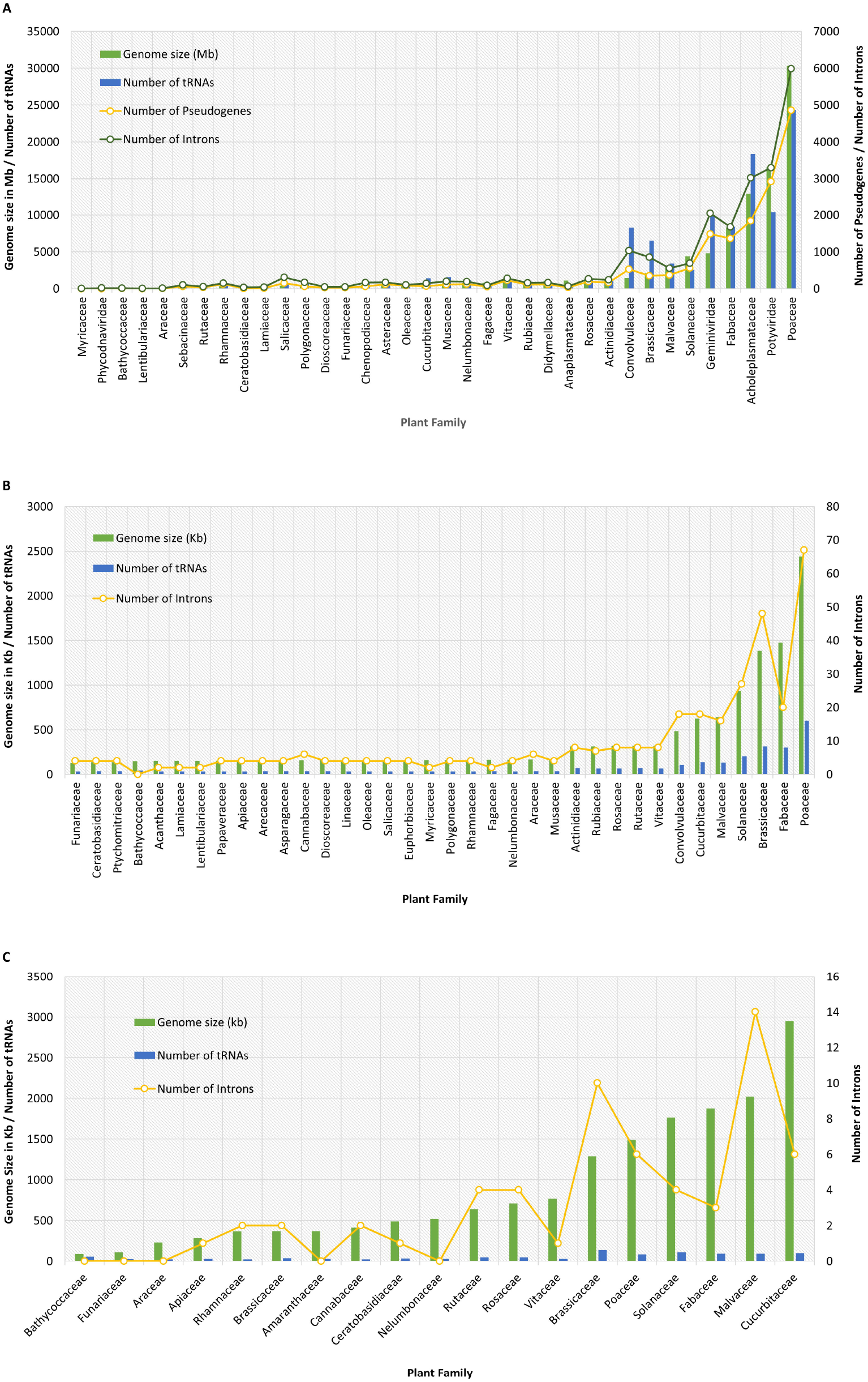
Relation between genome size, number of tRNAs, number of Pseudogenes and number of introns of nuclear, plastid and mitochondrial genomes. **A)** Shows relation between genome size in Mb, number of tRNAs, number of pseudogenes and number of introns in nuclear genome samples. **B)** Shows relation between genome size in Kb, number of tRNAs and number of introns in plastid genome samples. **C)** Shows relation between genome size in Kb, number of tRNAs and number of introns in mitochondrial samples.

Majority of the filtered tRNA sequences were from nuclear genome samples since, there were more tRNA filtration steps that were applied to the nuclear tRNA sequences. This can be evidently seen in the supplementary table 1 in which all the plants along with their initial tRNA count and post filtration tRNA sequence count are mentioned. In order to check the tRNA sequences of our database, we also cross validated our tRNA sequences with that of PlantRNA (10) database. Therefore, the all the tRNA sequences of PlantRNA database was retrieved and a database was developed using the ‘makeblastdb’ utility of the BLAST (9). All the true tRNA sequences of PtRNAdb database was provided as query input sequence data and ‘blastn’ was run using PlantRNA as the database. As a result of this analysis, we found 1346 tRNA sequences from the PlantRNA database that were exactly identical with 53,315 tRNA sequences of PtRNAdb database. Moreover, there were 1640 tRNA sequences from PlantRNA database with more than 75% identity with 90,022 tRNA sequences of PtRNAdb. Although, there is a huge amount of difference in the number of tRNA sequences reported in PlantRNA and PtRNAdb database, since, PlantRNA database reports tRNAs from only 12 plant species and PtRNAdb reports tRNAs from more than 100 plant species, but the evident number of tRNA matches indicate that, the data of PtRNAdb is reliable.

We have built covariance models using various utilities of infernal (11) like ‘cmbuild’, ‘cmcalibrate’ and ‘cmpress’ so that the users can structurally align their query sequences with different isotype and isodecoder models from various plants in PtRNAdb. This is available in the ‘ANALYZE’ module of PtRNAdb which is one of the key features of PtRNAdb. User will have to select a plant among one of the three genome levels followed by selecting the isoacceptors or isodecoder. The results of this module is available to download in two different formats i.e. in tabular format and in detailed output format exactly like shown in the result page of this module. To structurally align the user query sequence with the covariance model selected by the user, ‘cmscan’ utility of infernal is used. Since, Infernal’s cmscan or cmalign are always considered best for proper structural alignments, this module will provide the user with accurate alignments. Further, the consensus based study shows promising tRNA sequence consensus alignments and consensus tRNA structures. Although, some of the consensus structures of tRNA are not of very good quality and lack accuracy, but with future addition and curation of the existing data the consensus structures will improve and prove to be more reliable. The consensus study aims take into account whole groups of tRNAs based on isoacceptors and isodecoders, that is more accurate rather than, individual tRNA structures. However, the tRNA sequences which had only single isoacceptors or isodecoder do not have any consensus based alignments or structures. In such cases RNAfold which is a utility of Vienna RNA package (7) was used to build the tRNA structures and is shown on the detailed output page along with the RNA Dot Plot generated by RNAfold. This plant tRNA knowledgebase is aimed to provide large scale accurate information about tRNA biology and hope to aid in the experimental tRNA research.

## Future Directions

The PtRNAdb will be updated continuously with new and more accurate information along with better annotation of the existing tRNA sequences. We will continue to upgrade the quality of the web interface and offer new search possibilities. As the number of the tRNA entries increase in the database, we will provide file transfer protocol (ftp) and Application Programming Interface (API) options for the user to easily retrieve the data from PtRNAdb. A longer term aim of this work will be to enrich the biological information content of the databases, e.g. profiles, the description of tRNA gene expression profiles, the description of occurring tRFs and 3D structure models / clover leaf models of all the tRNAs.

## Acknowledgement

The authors are thankful to DBT (Department of Biotechnology)-eLibrary Consortium (DeLCON), India for providing access to e-resources. Authors are also thankful to Distributed Information Sub-Centre (Sub-DIC) of Department of Biotechnology (DBT) at NIPGR. The authors declare no competing financial interests.

## Author Contributions

D.D. and A.S. performed the data analysis work for the tRNA sequences of PtRNAdb. D.D. developed the most of the modules and web interface of the database including the Search, Browse, Blast and Analyze modules and A.S. developed the tRNA prediction module. D.D., S.Z. and S.K. wrote the complete manuscript. D.D. made all the figures and performed complete analysis that was included in the manuscript. S.K. conceived the idea and coordinated the project. S.K. agrees to serve as the author responsible for contact and ensures communication.

## Competing interests

The authors declare no competing interests.

## Table Legends

**Supplementary table 1:** Number of initial and post filtration tRNA sequences in Nuclear, plastid and mitochondrial genome levels.

## Notes

http://www.nipgr.ac.in/PtRNAdb/

### Competing Interest Statement

The authors have declared no competing interest.

## References

1. Lowe, T.M. and Eddy, S.R. (1997) tRNAscan-SE: a program for improved detection of transfer RNA genes in genomic sequence. Nucleic Acids Res., 25, 955–64.

2. Lowe, T.M. and Chan, P.P. (2016) tRNAscan-SE On-line: integrating search and context for analysis of transfer RNA genes. Nucleic Acids Res., 44, W54–7.

3. Bernhart, S.H., Hofacker, I.L., Will, S., Gruber, A.R. and Stadler, P.F. (2008) RNAalifold: improved consensus structure prediction for RNA alignments. BMC Bioinformatics, 9, 474.

4. Will, S., Joshi, T., Hofacker, I.L., Stadler, P.F. and Backofen, R. (2012) LocARNA-P: accurate boundary prediction and improved detection of structural RNAs. RNA, 18, 900– 14.

5. Thompson, J.D., Higgins, D.G. and Gibson, T.J. (1994) CLUSTAL W: Improving the sensitivity of progressive multiple sequence alignment through sequence weighting, position-specific gap penalties and weight matrix choice. Nucleic Acids Res., 22, 4673– 4680.

6. Huerta-Cepas, J., Dopazo, J. and Gabaldón, T. (2010) ETE: a python Environment for Tree Exploration. BMC Bioinformatics, 11, 24.

7. Lorenz, R., Bernhart, S.H., Höner zu Siederdissen, C., Tafer, H., Flamm, C., Stadler, P.F. and Hofacker, I.L. (2011) ViennaRNA Package 2.0. Algorithms Mol. Biol., 6.

8. Nawrocki, E.P. and Eddy, S.R. (2013) Infernal 1.1: 100-fold faster RNA homology searches. Bioinformatics, 29, 2933–5.

9. Altschul, S.F., Gish, W., Miller, W., Myers, E.W. and Lipman, D.J. (1990) Basic local alignment search tool. J. Mol. Biol., 215, 403–410.

10. Cognat, V., Pawlak, G., Duchêne, A.M., Daujat, M., Gigant, A., Salinas, T., Michaud, M., Gutmann, B., Giegé, P., Gobert, A., et al. (2013) PlantRNA, a database for tRNAs of photosynthetic eukaryotes. Nucleic Acids Res., 41.

11. Nawrocki, E.P. and Eddy, S.R. (2013) Infernal 1.1: 100-fold faster RNA homology searches. Bioinformatics, 29, 2933–2935.

